# Establishment of an eHAP1 Human Haploid Cell Line Hybrid Reference Genome Assembled from Short and Long Reads

**DOI:** 10.1101/822593

**Authors:** William D. Law, René L. Warren, Andrew S. McCallion

## Abstract

**Background:** Haploid cell lines are a valuable research tool with broad applicability for genetic assays. As such the fully haploid human cell line, eHAP1, has been used in a wide array of studies. However, the absence of a corresponding reference genome sequence for this cell line has limited the potential for more widespread applications to experiments dependent on available sequence, like capture-clone methodologies.

**Results:** We generated ~15x coverage Nanopore long reads from ten GridION flowcells. We utilized this data to assemble a *de novo* draft genome using minimap and miniasm and subsequently polished using Racon. This assembly was further polished using previously generated, low-coverage, Illumina short reads with Pilon and ntEdit. This resulted in a hybrid eHAP1 assembly with >90% complete BUSCO scores. We further assessed the eHAP1 long read data for structural variants using Sniffles and identify a variety of rearrangements, including a previously established Philadelphia translocation. Finally, we demonstrate how some of these variants overlap open chromatin regions, potentially impacting regulatory regions.

**Conclusions:** By integrating both long and short reads, we generated a high-quality reference assembly for eHAP1 cells. We identify structural variants using long reads, including some that may impact putative regulatory elements. The union of long and short reads demonstrates the utility in combining sequencing platforms to generate a high-quality reference genome *de novo* solely from low coverage data. We expect the resulting eHAP1 genome assembly to provide a useful resource to enable novel experimental applications in this important model cell line.

## Introduction

The vast majority of eukaryotic cells are diploid and many cellular models used experimentally are either diploid or polyploid. The presence of additional alleles, while evolutionarily beneficial, can pose challenges to genetic assays assessing loss of function mutations. This can occur through masking effects of a recessive mutation, or complicating experiments because of the necessity of retargeting unmodified alleles. To alleviate these challenges, haploid cell lines have been developed from a variety of species including medaka[1], rat[2], mouse[3, 4], and monkey[5]. In humans, a near haploid cell line containing the Philadelphia translocation, KBM-7, spontaneously arose from a subculture of a human leukemia tumor[6], although it remained diploid for chromosomes 8 and a portion of 15. Further work with these cells, in an unsuccessful attempt to induce pluripotency, resulted in a new cell line, termed HAP1; this line grew adherently and had lost a copy of chromosome 8[7]. Karyotyping of these cells also revealed loss of the Y chromosome. Finally, through the use of CRISPR/Cas9, HAP1 cells were genetically engineered to delete the diploid portion of chromosome 15, resulting in a fully haploid cell line termed eHAP1[8].

The HAP1 and eHAP1 cells have been used in a variety of experiments including drug screens[9, 10], host-virus interactions[7, 11–13], and genetic screens[14–16]. Despite the wide utility of these cells, only low coverage Illumina short read sequencing data has been generated for eHAP1 cells[8], resulting in challenges to producing a reference genome specific to this cell line. The generation of a more contiguous and generally higher quality reference genome would enable additional experimental uses such as sequence target capture, where knowledge of the underlying variants is critical. To this end, we employed the Oxford Nanopore Technologies Ltd. (ONT) GridION sequencing technologies to leverage the ability of long reads to uncover structural variants, which are difficult to detect using short reads alone. Additionally, combining long, but error prone read information with short, but highly accurate reads yields a more complete genome assembly than can be achieved using either technology individually[17–19].

## Results

### Generating a hybrid reference genome assembly of eHAP1 cells

Three independent replicates of high molecular weight genomic DNA was isolated from eHAP1s and prepared for Nanopore sequencing (Methods). We generated a total of ten individual libraries yielding 5 million reads and 48.1 Gb of sequence with an N50 of 31 Kb (Additional Table 1)[20]. The combined reads were aligned against the hg19 genome demonstrating an average coverage of ~15.6x (Fig 1a; Additional Figure 1).

**Figure 1:**
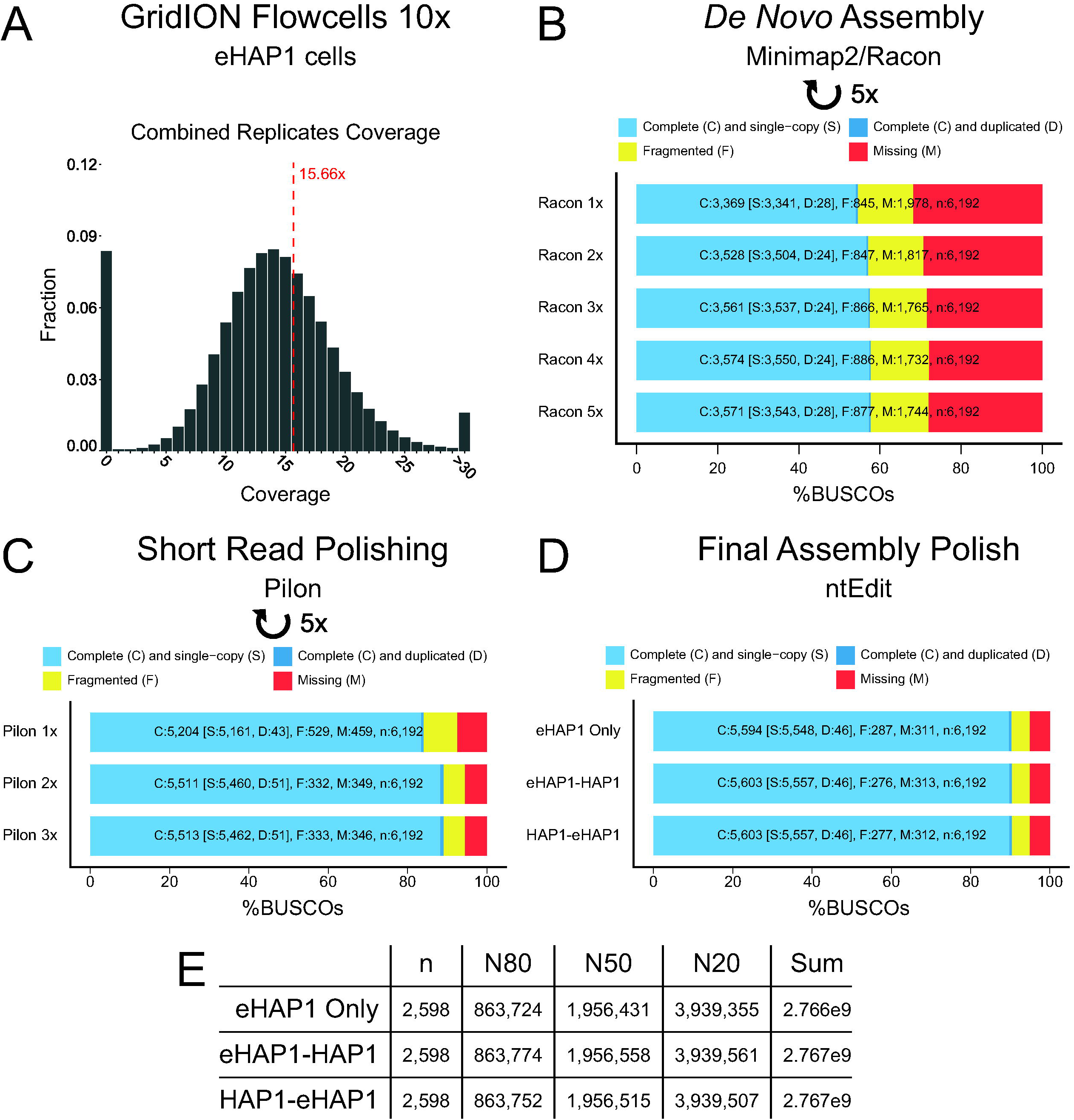
Polishing the eHAP1 reference genome. (**A**) Histogram of the combined ten ONT GridION flow cells coverage relative to the human genome (hg19). Greater than 30 reads were collapsed into a single bin, and the red line indicates the average mean coverage. BUSCO[24, 25] scores were calculated after five rounds of Racon[23] polishing (**B**) and three rounds of Pilon[26] (**C**). Left indicates the number of rounds of each program, and bars display BUSCO notation. (**D**) ntEdit[27] was performed using the eHAP1 short reads on the 5x Racon/3x Pilon (eHAP1 Only) polished assembly and using eHAP1 then HAP1 (eHAP1-HAP1) or HAP1 then eHAP1 (HAP1-eHAP1) short reads. BUSCO sores were calculated after each round. (**E**) ABySS [28] contiguity statistics were calculated for the three ntEdit polished assemblies.

Next, the reads were aligned against themselves to generate a *de novo* reference assembly. Subsequently, error correction was performed using five successive iterations of minimap2[21], miniasm[22], and Racon[23] (Fig 1b). After each round, we calculated the BUSCO (Benchmarking universal single-copy orthologs) scores[24, 25] to evaluate the quality of the consensus assembly. We observed a 5.5% increase in complete BUSCOs from the first (3,369 complete BUSCOs) to the last (3,571 complete BUSCOs) round of error correction; however, after five rounds we observed no appreciable improvement using long reads alone.

To further improve the quality of the reference genome, we incorporated previously generated Illumina short reads[8] utilizing Pilon[26] to further polish the assembly (Fig 1c). While eHAP1 cells are a direct derivative of the E9 clone (~6x whole genome sequencing (WGS) coverage), we also chose to use short reads from a parallel eHAP1 clone (A11; ~6x WGS coverage) to obtain ~12x total coverage. To assess the quality of the reference genome, we again used BUSCO scores and saw a large improvement with just one round of Pilon polishing. The number of complete BUSCOs increased markedly from 3,571 to 5,204 (31.4%) and the number of fragmented or missing decreased by 1,633. Using the polished output from Pilon, we repeated the short read polishing two additional times and observed moderate improvements in BUSCO scores. Finally, due to the low Illumina sequencing coverage, we employed an additional polishing step utilizing ntEdit[27], which functions well in low sequence coverage situations. We observed a slight improvement recovering an additional 81 complete BUSCOs, ultimately obtaining 90% BUSCO completeness with ~5% listed as fragmented or missing, respectively. Overall, we were able to generate a high quality *de novo* reference genome using a combination of low coverage Nanopore long reads and Illumina short reads.

Initially, we did not include the ~20x WGS coverage from the parental HAP1 cells to avoid any potential diploid regions or eHAP1 specific sequence variants. However, to assess if additional sequencing depth could further improve the reference genome, by resolving indels causing frameshift errors, we polished the reference genome using the parental HAP1 data. Starting with the eHAP1 reference assembly polished by 5x Racon and 3x Pilon, we used ntEdit to polish the assembly using the eHAP1 data first, followed by a second round using the HAP1 data, or the reverse order. In either case, we saw virtually no improvement in BUSCO scores, nine additional complete BUSCOs, compared with using the eHAP1 data alone (Fig 1d). We also utilized ABySS[28] to assess the assembly contiguity statistics of the three polished references (Fig 1e), and observe similar statistics across the three. To prevent potentially confounding variants from the parental cell line data, we focused on the polished reference assembly using eHAP1 reads only. By combining long and short reads, we were able to assemble a high-quality reference genome for the eHAP1 cells.

### Long reads identify multiple structural variants

We next sought to identify structural variants (SVs) present in the eHAP1 cells. To do this, we used the structural variant caller Sniffles[29]. We first aligned the Nanopore reads to the hg19 reference genome using NGMLR[29] and passed the output into Sniffles to identify structural variants. Using the default threshold of a minimum of ten reads supporting the SV, we identified 11,451 SVs (Fig 2a; Additional Table 2); however, due to the lower coverage, we reduced the minimum number of reads to five, which yields 18,295 SVs (Additional Table 3). Despite this, the SV types between the two sets are very similar with a vast majority of SV subtypes identified as either deletions (Fig 2b) or insertions (Fig 2c). Additionally, using the five read threshold, many of the deletions (6,268/8,665; ~72.3%) or insertions (5,187/7,683; ~67.5%) are small, less than 250 base pairs (bp). Critically, ONT reads are known to have biases in deletions potentially due to difficulty in basecalling [29, 30]. We find 1,306 (15.1%) deletions detected by Sniffles contain homopolymeric runs of at least 20 bp and an additional 954 (11%) deletions overlapping dinucleotide repeats of at least 10 bp. This implies these detected deletions may be a technical artifact, rather than genuine rearrangements.

**Figure 2:**
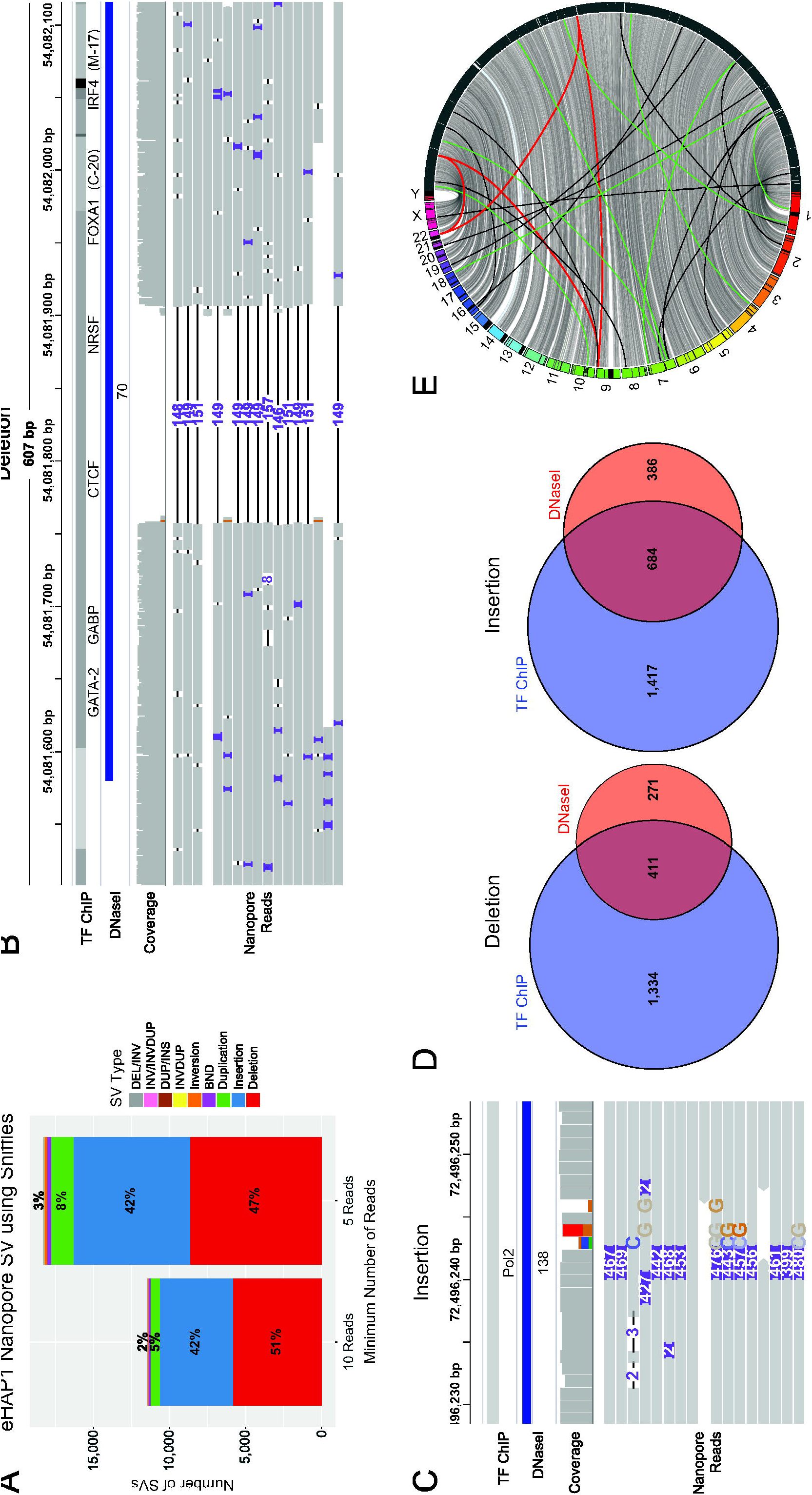
Structural variant analysis of the eHAP1 cell line. (**A**) A breakdown of the types of structural variants (SV) identified by Sniffles[29] from the eHAP1 cell line. The X-axis refers to the minimum number of reads required to support a SV. Visualization using IGV genome browser[42] of a deletion (**B**) or insertion (**C**) DNaseI hypersensitivity sites (DNaseI) [31, 32] and transcription factor binding sites (TF ChIP) [31, 33]. The numbers inside the SV indicate the size in base pairs. (**D**) Venn diagram illustrating the number of SVs overlapping TF ChIP or DNaseI sites. (**E**) A Jupiter plot[34] of the eHAP1 only polished assembly against the human genome (GRCh38). GRCh38 chromosomes are displayed incrementally from 1 (bottom, red) to Y (top, fuchsia) on the left while scaffolds (grey with black outlines) are displayed on the right side of the rim. The highlighted lines indicate potential translocations. The black lines indicate potential translocations not found using Sniffles. The green lines indicate potential chromosomal translocations where Sniffles also indicates a translocation between the two chromosomes. The red lines indicate the Philadelphia translocations, identified here and by Sniffles.

In spite of the potential false positives, once the SVs overlapping repeats are removed, we identify 271 deletions and 386 insertions that specifically overlap an open chromatin region[31, 32] (OCR; Fig 2d) identified by ENCODE in any assessed cell line. There are an additional 1,334 deletions and 1,417 insertions that do not overlap an OCR but do impact a transcription factor binding site (TFBS) [31, 33]. Finally, there were 411 and 684 regions that disrupt both an OCR and TFBS for insertions and deletions, respectively. Collectively, we find a large number of SVs impacting putative regulatory regions that could impact experimental design or interpretations.

Other types of SVs identified include translocations. As the parental KBM-7 cell line contained the Philadelphia chromosome that was retained throughout subcloning, we anticipated identifying a translocation between chromosomes 9 and 22. Sniffles correctly identified the translocation (chr9: 133,681,711 - chr22:23,632,359; hg19) directly within in the *BCR* and *ABL* genes. In all, we detect 250 translocations, of which 60 are classified as precise, indicating confidence of the exact breakpoint position at the nucleotide level. Interestingly, Sniffles detects 31 translocations involving the Y chromosome, despite karyotyping data suggesting it was lost between the KBM-7[6] to HAP1[7]. Our polished assembly potentially suggests a translocation of the Y chromosome onto the X chromosome, but the alignment quality to the Y is moderate (~60%). While Sniffles suggests the Y chromosome may have been broken and scattered throughout the genome, other data indicates it may have been lost entirely, and additional experiments are necessary to distinguish these possibilities.

Finally, we generated an assembly consistency (Jupiter) plot[34] showing the polished eHAP1 reference genome scaffolds against GRCh38 (Fig 2e). From it, we clearly identify the Philadelphia translocation and additionally identify other SVs that corroborate Sniffles’ findings in the ONT reads. Utilizing long reads exclusively, we were able to discover a variety of SVs, many of which are insertions or deletions potentially impacting OCRs, as well as larger translocations.

## Discussion

We employed ONT long read sequencing to improve the reference genome quality and identify SVs of an important human cell line, eHAP1. By utilizing previously published Illumina short read data with low coverage (~12x) and combining it with our long read data, we were able to generate a high-quality hybrid genome assembly with complete BUSCO scores of 90%. This required the use of a variety of polishing tools including Racon[23], Pilon[26], and ntEdit[27].
While we observed the greatest improvement through one round of Pilon utilizing short reads, it is important to note it required the greatest amount of computational resources per round: 3-4 days, 48 processors, and 384 Gb of RAM. In comparison, ntEdit required 36 minutes, 48 processors, and 22.2 Gb of RAM. While ntEdit benefited from a Pilon polished reference, we did not see any appreciable resource reduction between sequential Pilon rounds. Additionally, we did not need to employ ntEdit prior to Pilon polishing, but it may be beneficial in situations where computational resources are limited. Regardless of the computational requirements, Pilon produced the largest increase in BUSCO scores despite the relatively low sequencing coverage of Illumina data. Providing additional accurate short reads, using either deeper coverage Illumina sequencing or linked reads, would likely improve base pair accuracy, scaffolding, and contiguity of the reference generated here; however, utilizing the deeper (~20x) coverage of the parental HAP1 cell line[8] had little impact on the final quality of the reference genome generated, as assessed by BUSCO analysis.

One of the greatest advantages of long reads is the ability to easily detect structural rearrangements. We were able to use the Nanopore data aligned against the human reference genome to identify over 18,000 SVs, a majority of which were small insertions or deletions. It is important to note that Nanopore reads are prone to over-calling deletions residing in repetitive regions of the genome. While a portion of these deletions may not be validated, Sniffles was able to find highly confident insertions and deletions, some of which reside within OCR and within TFBS. This type of information would be useful for experiments interested in using this cell line to assess regulatory regions[35] or in cases where the eHAP1 cells are used as primary genomic DNA isolation for capture-clone experiments[36, 37]. Additionally, Sniffles detected the presence of the Philadelphia chromosome translocation with high confidence using the long reads exclusively.

In summary, we applied Nanopore long read sequencing technology to an important human haploid cellular model. Utilizing a combination of long and short reads, we were able to generate a high-quality reference genome and demonstrate the utility of a hybrid assembly despite comparatively low sequencing coverage. We anticipate this work will enable novel applications of eHAP1 cells, such as capture sequencing experiments and targeted CRISPR screens, to be conducted in an accelerated time frame.

## Methods

### eHAP1 cell culture

eHAP1 cells were purchased from Horizon Discovery (SKU: c669). The cells were cultured using the following growth media: 445 mL IMDM media (Gibco: 12440-053), 50 mL FBS, and 5 mL 100x Pen/Strep. Cells were passaged every 2-3 days at a ratio of 1:5. The cells were rapidly expanded post purchase to reduce the number of passages and possible ploidy changes, prior to genomic DNA isolation.

### Genomic DNA isolation, library prep, and sequencing

Genomic DNA was harvested from 5 million cells using the Circulomics Nanobind CBB Big DNA kit (Part #NB-900-001-01). The DNA was extracted following the included handbook (v1.7) protocol for “Cultured Mammalian Cells – HMW” with minor modifications. Specifically, cells were vortexed intensively (1 second pulses, 10x pulses), the final DNA was pipetted 10 times through a p200 tip, and immediately prior to library preparation, the DNA was run through a 28G needle five times. This was done to help the DNA into solution with minimal effect on length.

The genomic DNA was prepared using the Nanopore Ligation Sequencing Kit (SQK-LSK109) following the manufacturer’s protocol (GDE_9063_v109_revD_23May2018). An initial starting amount of 1 μg of genomic DNA was used, and after library preparation, a final amount of 250-400 ng was obtained. Regardless of the final mass of DNA obtained, the entire library preparation was subjected to R9.4 flowcells (FLO-MIN106) in a 1:1 library preparation:flowcell ratio. Basecalling was performed using guppy v2.3.7. Run statistics were calculated using NanoPlot (-t 12) [20].

### De novo genome assembly and polishing

The Nanopore long read .fastq files from all 10 replicates were combined and aligned against each other using minimap2 (v2.16-r922)[21] with the -x ava-ont and -t 24 flags. A layout was generated using miniasm (v0.3-r179)[22] with the -t 24 flag, and the resulting .gfa file was converted into a .fasta format using awk. Then, the original Nanopore reads were aligned against the .fasta file using minimap2 (-t 24), and the resulting pairwise mapping format (.paf) file was combined with the .fasta and the combined .fastq files to be polished using Racon (v1.3.3)[23] (-t 24). The output, a .fasta polished formatted file, was passed back to minimap2, and the reads were aligned a second time. This process was repeated a total of five times.

After five rounds, the resulting polished .fasta file was used as a reference to map the previously generated[8] Illumina short reads using minimap2 (-ax sr). Data from both clone E9 and A11 were mapped independently, and the resulting .sam files were converted to sorted and indexed .bam files using samtools (v1.9)[38]. Finally, both .bam files were used to polish the .fasta file using Pilon[26] (-xmx700G; v1.22). The resulting Pilon polished .fasta format was used to re-map the short reads using minimap2. This process was performed a total of three times.

The resulting Pilon-polished assembly was used as input for ntEdit (v1.2.2)[27]. Briefly, we ran ntEdit iteratively 3 times (-k 50-40 step 5, -i 5 -d 5 -m 1 -t 48) each with k=50, k=45 and k=40 kmer Bloom filters derived from running ntHits (v0.0.1 --outbloom --solid -b 36 -k 50-40 step 5 -t 48) on the combined Illumina short read data. Run time and memory usage was benchmarked on a CentOS 7 system with 128 Intel(R) Xeon(R) E7-8867 v3 CPUs @ 2.50GHz.

After each round of polishing, regardless of the program, BUSCO (v3.1.0)[24, 25] scores were assessed. The program was run in --mode genome, with --cpu 24 and --blast_single_core. The files were compared against the euarchontoglires_odb9 lineage. ABySS (v2.1.0) [28] statistics were calculated using the abyss-fac function.

### Structural variant detection

Structural variants were detected using Sniffles[29] after alignment of the long reads against the hg19 reference genome using NGMLR[29] (-t 24, -x ont). The resulting .sam file was converted into a sorted .bam using samtools[38] and passed onto Sniffles in either default mode (-s 10) or five read minimum (-s 5). The resulting .vcf file was parsed into SV types using grep, and figures were made using ggplot2[39].

The overlap with ENCODE DNaseI [31, 32] and TF ChIP [31, 33] datasets was performed using the UCSC Table Browser[40]. A .bed file was made from structural variants detected (five read minimum) using the left-most coordinate and adding one basepair. The .bed file was uploaded to the UCSC Table Browser and intersected, with 100% overlap, with the “wgEncodeRegDnaseClusteredV3”[31, 32] or “encRegTfbsClustered”[31, 33]. The resulting regions were filtered into insertions and deletions using Sniffles SVTYPE information, and the sequences were further filtered for repeats. For homopolymeric repeats, insertions or deletions containing 20 identical basepairs in a row and dinucleotide repeats of 10 pairs of any two basepairs consecutively were removed.

### Jupiter plot generation

An assembly consistency (Jupiter v1.0) plot[34] of the polished reference eHAP1 genome was generated. As part of the Jupiter plot pipeline, scaftigs from the largest eHAP1 scaffolds, consisting of 75% (NG75) of the genome, were aligned to GRCh38 with minimap2 (v2.17-r941) and plotted with Circos (v0.69-6_1)[41].

## Data Accession

The raw Nanopore reads generated in this study are available upon request. The previously generated[8] Illumina short reads for clone A11 (SRR1518295) and E9 (SRR1518293) are available from the NCBI Sequence Read Archive. The final polished .fasta formatted eHAP1 reference genome is available upon request. The eHAP1 cell line may be purchased from Horizon Discovery. All custom programs and intermediate files are available upon request.

## Abbreviations

SV: Structural Variant
BUSCO: Benchmarking Universal Single-Copy Orthologs
bp: base pairs
WGS: Whole Genome Sequencing
OCR: Open chromatin region
ONT: Oxford Nanopore Technologies Ltd
TFBS: Transcription Factor Binding Site

## Acknowledgments

This work was supported from the NIH (MH106522) and the National Institutes of Health [2R01HG007182-04A1]. This work was also supported through internal funding from the Johns Hopkins University School of Medicine as part of the Core Coins Program. We acknowledge assistance for Nanopore sequencing from the Genetic Resources Core Facility High-Throughput Sequencing Core. The content of this work is solely the responsibility of the authors, and does not necessarily represent the official views of the National Institutes of Health or other funding organizations.

We thank David Mohr for providing guidance and computational resources, Jeffrey Burke from Circulomics for assistance in genomic DNA extraction, and Paul W. Hook and Sarah A. McClymont for critical reading of the manuscript. Additional computational resources were provided by the Maryland Advanced Research Computing Center (MARCC).

---

**Additional Figure 1:**
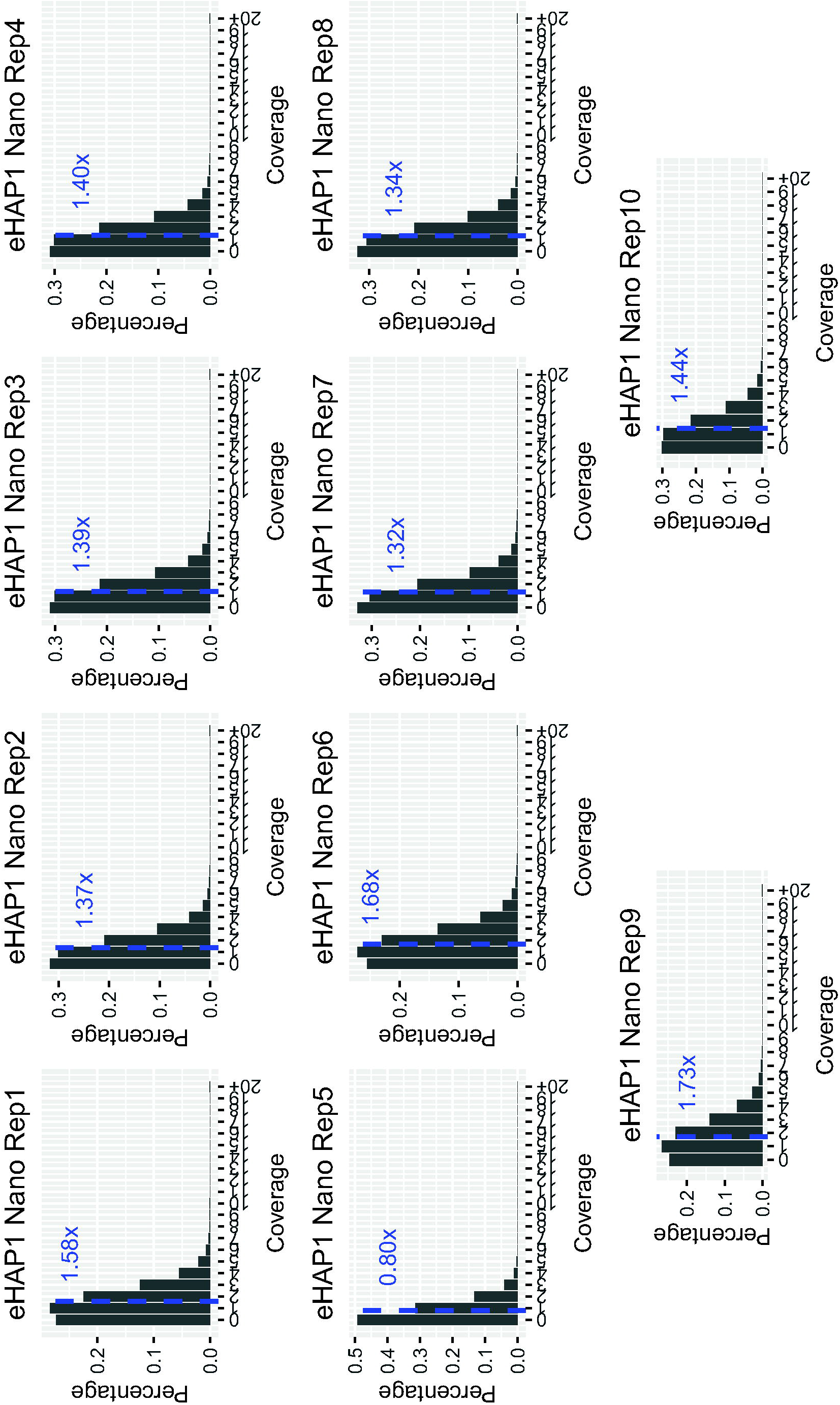
Coverage per individual flow cell. Histogram of the coverage plots for each of the ten replicates. The blue line and number indicates the mean coverage for a flow cell. Coverages above 20 were collapsed into a single bin (20+).

**Additional Table 1**: <u>*NanoPlot statistics per individual flow cell*</u>. NanoPlot[20] statistics were computed for each flow cell. For appropriate statistics, the mean was calculated across the ten replicates (column L). NanoPlot was also run on the combined flow cells (column M).

**Additional Table 2**: <u>*Structural variants detected by Sniffles (Default; ten reads)*</u>. Sniffles was run in default mode, ten reads minimum supporting SV calls, using the long reads generated from eHAP1 cells. Column information is indicated in the header, and additional information can be found on the Sniffles wiki: https://github.com/fritzsedlazeck/Sniffles/wiki/Output.

**Additional Table 3**: <u>*Structural variants detected by Sniffles (Default; five reads)*</u>. Sniffles was run in using five read minimum supporting SV calls, using the long reads generated from eHAP1 cells. Column information is indicated in the header, and additional information can be found on the Sniffles wiki: https://github.com/fritzsedlazeck/Sniffles/wiki/Output.

